# The unforeseen intracellular lifestyle of *Enterococcus faecalis* in hepatocytes

**DOI:** 10.1101/2021.09.30.462169

**Authors:** Natalia Nunez, Aurélie Derré-Bobillot, Goran Lakisic, Alexandre Lecomte, Françoise Mercier-Nomé, Anne-Marie Cassard, Hélène Bierne, Pascale Serror, Cristel Archambaud

**Affiliations:** Université Paris-Saclay, INRAE, AgroParisTech, Micalis Institute, 78350, Jouy-en-Josas, France; Université Paris-Saclay, INSERM, CNRS, Institut Paris Saclay d’Innovation Thérapeutique, Châtenay-Malabry, France; Université Paris-Saclay, INSERM U996, Inflammation, Microbiome and Immunosurveillance, 92140, Clamart, France

**Keywords:** *Enterococcus faecalis*, pathobiont, hepatocytes, intracellular lifestyle, liver

## Abstract

*Enterococcus faecalis* is a bacterial species present at a sub-dominant level in the human gut microbiota. This commensal turns into an opportunistic pathogen under specific conditions involving dysbiosis and host immune deficiency. *E. faecalis* is also the only intestinal pathobiont identified to date as contributing to liver damage in alcoholic liver disease. We have previously observed that *E. faecalis* is internalized in hepatocytes. Here, the survival and fate of *E. faecalis* was examined in hepatocytes, the main epithelial cell type in the liver. Although referred to as an extracellular pathogen, we demonstrate that *E. faecalis* is able to survive and divide in hepatocytes, and form intracellular clusters in two distinct hepatocyte cell lines, in primary mouse hepatocytes, as well as *in vivo*. This novel process extends to kidney cells. Unravelling the intracellular lifestyle of *E. faecalis*, our findings contribute to the understanding of pathobiont-driven diseases.

## Introduction

Among chronic liver diseases, alcoholic liver diseases, non-alcoholic fatty liver diseases, chronic viral hepatitis, and hemochromatosis are the most common worldwide diseases ^1^. These liver disorders are associated with prolonged alcohol consumption, infections, autoimmune diseases, and genetic and metabolic disorders. Recently, dysbiosis within the intestinal microbiota, associated with a decrease in the diversity of microbial populations and the proliferation of potentially pathogenic species, has been recognized as an important additional factor in the etiology of liver diseases ^2^. *Enterococcus faecalis* is a sub-dominant commensal bacterium of the human gut microbiota and can become pathogenic under specific conditions involving gut dysbiosis and host immune deficiency ^3^. While antibiotic treatments are well known to cause enterococcal overgrowth, which may lead to systemic infection in immunocompromised patients, other drug treatments trigger intestinal dysbiosis. For example, the long-term use of proton pump inhibitors (PPIs), frequently prescribed in patients with liver diseases, is associated with harsh effects such as the development of spontaneous bacterial peritonitis and an increased risk of developing hepatic pyogenic abscesses ^4, 5^. PPI treatments are associated with significant changes in the intestinal microbiota, including an increase in the genera *Enterococcus, Streptococcus*, and *Staphylococcus* and the species *Escherichia coli* ^6^. Notably, patients with liver disease frequently present a dysbiotic microbiota with an overgrowth of enterococci ^7, 8^.

While a link between vancomycin-resistant enterococci (VRE) intestinal domination and bloodstream infections has been reported ^9-11^, the translocation of enterococci to the liver has not yet been fully established in patients. In contrast, we and others reported enterococcal translocation from the gut to the liver in rodent models ^12-15^. In alcohol-mediated liver disease, ethanol consumption increases intestinal permeability by disrupting the gut microbiota and tight-junction integrity. Alcohol and PPI treatment are known to benefit *E. faecalis* translocation, which promotes inflammation mediated by toll-like receptors (TLR) on Kupffer cells that recognize extracellular *E. faecalis* in the liver ^14^. Their findings were corroborated by the significant increase in *Enterococcus* in the stool of healthy individuals after two weeks of treatment with PPIs and in chronic alcohol users taking PPIs ^14^. More recently, it has been shown that the severity of alcoholic hepatitis and mortality of patients with alcoholic hepatitis are consistent with the presence of *E. faecalis* expressing cytolysin, a toxin capable of lysing bacteria and cells ^16^.

*E. faecalis* is generally described as an extracellular bacterium capable of entering and surviving in mammalian cells. *E. faecalis* can enter and survive in non-professional phagocytic cells, like intestinal epithelial cells, urothelial cells from the bladder, and endothelial cells ^17-22^. Several invasion pathways relying on cytoskeleton components have been proposed ^17, 22^. Upon internalization in epithelial cells, *E. faecalis* has been observed in endosomal compartments or organized into intracellular colonies ^17, 18, 23^. If enterococci survive within macrophages for extended periods, likely due to their ability to reduce host cell autophagy and to prevent its delivery in typical LC3^+^ autophagic compartments ^24-26^, how they survive and persist in epithelial cells remains to be established. Conversely, intestinal epithelial autophagy can be activated by *E. faecalis* and coincides with the formation of autophagosomes surrounding *E. faecalis* ^27^. The fate of *E. faecalis* once internalized in epithelial cells seems to be much more complex to assess and probably depends on the specialization of different epithelial cell types.

We previously observed that *E. faecalis* is internalized in HepG2 hepatic cells ^28^. Considering increasing evidence that intestinal *E. faecalis* may be able to reach the liver of patients, this study examined the interactions between *E. faecalis* and hepatocytes, which account for 70% of hepatic cells in the liver, in more detail. Using two human hepatocyte models of infection, *ex vivo* and *in vivo* models, the fate of *E. faecalis* was investigated after its internalization in hepatocytes.

## Results

### *Intracellular growth of* E. faecalis *during the infection of human hepatocytes*

To investigate the fate of *E. faecalis* in hepatocytes, the capacity of *E. faecalis* strain OG1RF, a human isolate, was assessed to determine its invasion and survival ability within human Huh7 hepatocytes by comparing the number of intracellular bacteria to the initial inoculum (Figure 1A, left inset). The hepatocytes were infected for 3 h with *E. faecalis* before the addition of a gentamicin- and vancomycin-containing medium to kill the extracellular bacteria. One h after antibiotic treatment, the median level of invasion of Huh7 by OG1RF was about 0.08%. Notably, the level of intracellular enterococci reached a median value of about 0.16% 24 h post-antibiotic treatment (pa) This indicates that the intracellular bacterial ratio doubled in 24 h. This percentage remained high 48 h pa, indicating that intracellular *E. faecalis* bacteria not only survive but proliferate within Huh7 cells. The antibiotic protection assay is commonly used to study intracellular pathogens, but several studies have reported that some antibiotics penetrate and accumulate within the host cells, affecting the intracellular growth of the pathogen ^29, 30^. The addition of amoxicillin, a well-known β-lactam antibiotic used to kill intracellular pathogens ^31^, was tested to determine its effect on the increase of intracellular enterococci during infection. As shown in Figure 1A (right insert), while the percentage of intracellular *E. faecalis* was comparable 1 h pa to that observed without amoxicillin, the percentage decreased dramatically at 24 and 48 h. These data indicate that the penetration and accumulation of amoxicillin in Huh7 cells decreases the intracellular level of enterococci in hepatocytes. As amoxicillin mainly targets dividing bacteria by blocking the cross-linking process during peptidoglycan synthesis, this result indicates that *E. faecalis* bacteria are able to divide within the host cell.

**Fig 1.**
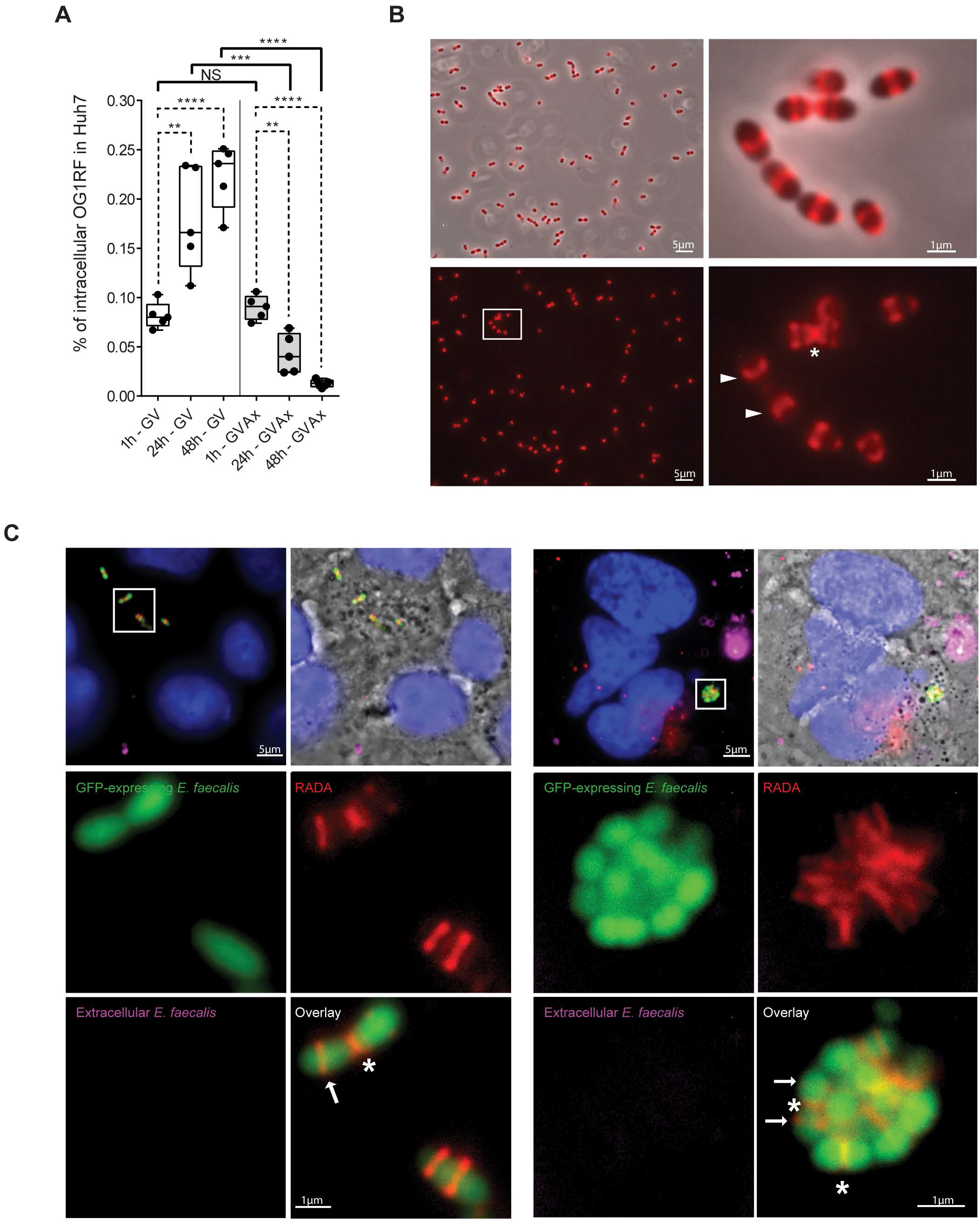
*Enterococcus faecalis* divides during the infection of human hepatocytes. Exponentially growing *E. faecalis* OG1RF cultures were used to infect Huh7 cells for 48 h. **(A)** The percentage of intracellular bacteria in Huh7 cells was determined as the ratio of intracellular bacteria of the initial inoculum and compared in two antibiotic cocktails containing gentamycin (G) and vancomycin (V) with or without amoxicillin (Ax). Data are represented by box-whisker plot (min to max) of five independent experiments. Each dot represents one independent experiment. The horizontal bar indicates the median value. Statistical analysis was performed using an unpaired Student’s t test. Asterisks indicate a p-value considered statistically significant (**, P < 0.01; ***, P < 0.001; ****, P < 0.0001), NS, non-significant difference. **(B)** *E. faecalis* growing in BHI-rich medium were labeled with the orange-red TAMRA-based fluorescent D-amino acid (RADA) labeling peptidoglycan in live bacteria for 40 min. The RADA signal was detected in the mid cell corresponding to the septal ring (asterisks) and to equatorial rings (white arrow heads). **(C)** Huh7 cells were infected with GFP-expressing *E. faecalis* for 36 h. Twelve hours after the addition of the antibiotic-containing medium, infected cells were incubated with RADA. Representative micrographs of individual *E. faecalis* **(left panels)** and enterococcal clusters **(right panels)** in Huh7 cells are shown from two independent experiments. The image is an overlay of the phase contrast, intracellular *E. faecalis* (green channel), extracellular *E. faecalis* (pink channel), and nuclei (blue channel). The scale bar corresponds to 5 µm. One framed enterococcal cluster is shown at a higher magnification below (Bar: 1 μm). The framed image is an overlay of intracellular *E. faecalis* (green channel) and RADA (red channel). Asterisks indicate signals detected in the mid cell corresponding to the septal ring. White arrows indicate equatorial ring signals.

To confirm this hypothesis, a fluorescent D-amino acid, which is incorporated into the bacterial cell wall and labels the newly formed peptidoglycan in live bacteria ^32^, was used. First, the RADA molecule was added to enterococcal exponential cultures and a fluorescent signal was detected in the mid cell corresponding to the septal ring and to equatorial rings (Figure 1B). Next, GFP-expressing *E. faecalis* infected Huh7 hepatocytes were incubated with RADA (orange-red TAMRA-based fluorescent D-amino acid) in the cell culture medium for 24 h, after 12 h in the antibiotic-containing medium. Intracellular *E. faecalis* showed an incorporation of the RADA molecule in their cell wall compared to the remaining antibiotic-killed extracellular bacteria (Figure 1C and Figure S1). Notably, RADA-labelled cocci and diplococci were organized into groups. For both, localization patterns of incorporated RADA included signals detected in the mid cell corresponding to the septal ring and to duplicated equatorial ring signals corresponding to the elongation step of cell division, similar to those obtained from *E. faecalis* growing in rich medium. RADA incorporation definitively supports intracellular growth during *E. faecalis* infection of hepatocytes.

### *Formation of enterococcal clusters accompanies* E. faecalis *growth within hepatocytes*

To get insights into the *E. faecalis* growth in hepatocytes, differential immunofluorescence labelling was performed to track intracellular *E. faecalis* internalized in Huh7 hepatocytes ^33, 34^. The presence of intracellular cocci and diplococci organized in chains or groups, which we hereafter called “clusters” when the number of bacteria inside included at least four cocci, were confirmed (Figure 2A). The appearance of these intracellular clusters was quantified in Huh7 hepatocytes between 30 min and 48 h pa, and the numbers of cocci per cluster was determined. Enterococcal clusters were detectable after 30 min in 3% of the infected cells. Compared to 30 min, the median value of the Huh7 cells exhibiting at least one enterococcal cluster significantly increased to 9% after 24 h pa (Figure S2A). The number of cocci per cluster showed a significant increase in the size of the intracellular enterococcal clusters in the Huh7 hepatocytes during infection. While the median value of the number of enterococci within a cluster was four at 30 min pa, it reached seven bacteria 48 h later (Figure S2B). Notably, clusters including more than ten bacteria were rare at 2 h, whereas those with ten and more than 20 cocci increased between 2 and 48 h pa (Figure 2B). These results revealed the formation of enterococcal clusters whose size increased during infection. The formation of *E. faecalis* OG1RF clusters in two other cell types, HepG2 cells and in primary mouse hepatocytes, was also examined (Figure 2C). At 24 h pa, intracellular clusters were observed in both cell types, as described in Huh7 cells. Detection of clusters in primary mouse hepatocytes showed that the formation of intracellular enterococcal clusters was independent of the immortalized phenotype of the two hepatocyte cell lines. Together, our data show that intracellular *E. faecalis* growth is accompanied by the formation of clusters during hepatocyte infection.

**Fig 2.**
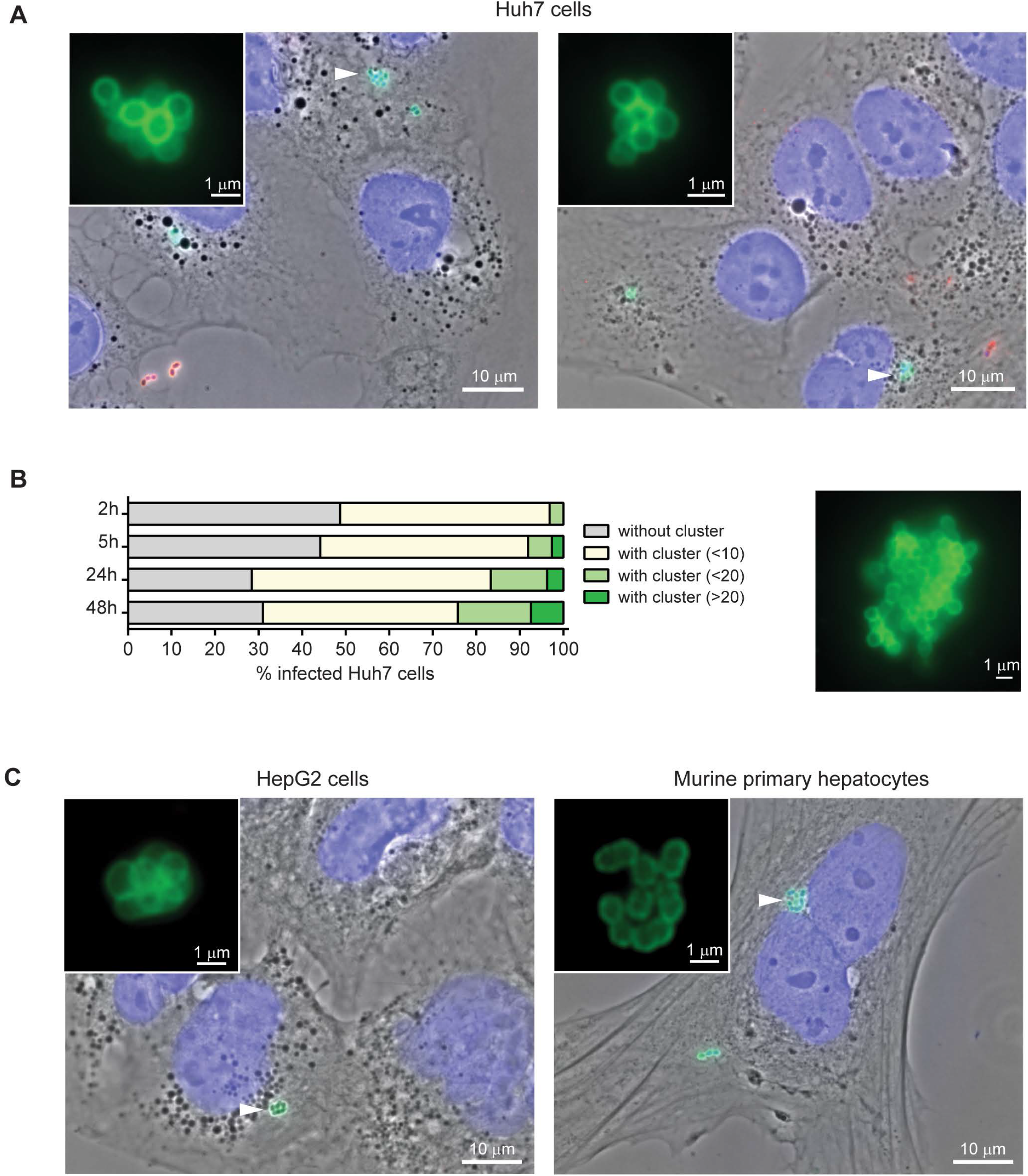
Intracellular *E. faecalis* forms clusters within hepatocytes. **(A)** Representative micrograph of Huh7 cells infected for 5 **(left panel)** and 48 h **(right panel)** with *E. faecalis* OG1RF observed in six independent experiments (objective 100x). The image is an overlay of the phase contrast, intracellular *E. faecalis* (green channel), antibiotic-killed extracellular *E. faecalis* (red channel), and nuclei (blue channel). White arrowheads indicate intracellular clusters. The scale bar corresponds to 10 µm. Two framed enterococcal clusters are shown at a higher magnification (Bar: 1 μm). **(B)** Quantification of the percentage of cells according to the number of enterococci within the intracellular cluster in Huh7 cells. For each time point, at least 3,400 cells were examined at low magnification (objective 40×) from three independent experiments. A framed enterococcal cluster exhibiting more than 20 cocci is shown at a higher magnification (Bar: 1 µm). **(C)** Representative micrograph of HepG2 and primary mouse hepatocytes infected 24 h with *E. faecalis* OG1RF observed in two independent experiments. The image is an overlay of the phase contrast, intracellular *E. faecalis* (green channel), extracellular *E. faecalis* (red channel), and nuclei (blue channel). The scale bar corresponds to 10 µm. For each cell type, one intracellular enterococcal cluster indicated by a white arrowhead is shown at a higher magnification (Bar: 1 μm).

### Intracellular enterococcal clusters form in the mouse liver, and enterococcal infection associates with sequential changes in Kupffer macrophages and neutrophil populations

To track enterococcal clusters within hepatocytes *in vivo, E. faecalis* strain OG1RF, expressing the *luxABCDE*(*lux*) operon from *Photorhabdus luminescens* driven by a constitutive promoter, was used. Compared to non-infected control mice, a luminescent signal was detected 6 h post-infection (pi) in mice infected intravenously (Figure S3). The signal emitted by the liver remained mostly stable or increased 24 h pi. Since the border of the bigger left lobe emitted a very strong signal (Figure S3), an immunohistological analysis was performed on this delimitated area. As shown in Figure 3A, the presence of *E. faecalis* on lobe liver sections was detected through green signals emitted from round foci compared to noninfected mice. The size of these *E. faecalis* infection foci increased at 24 h pi (Figure 3A). At higher magnification, enterococcal clusters were clearly identified within multinucleated hepatocytes, highly expressing the protein claudin-1 ^35^, indicating that the intracellular *E. faecalis* division in hepatocytes also occurs *in vivo* (Figure 3B).

**Fig 3.**
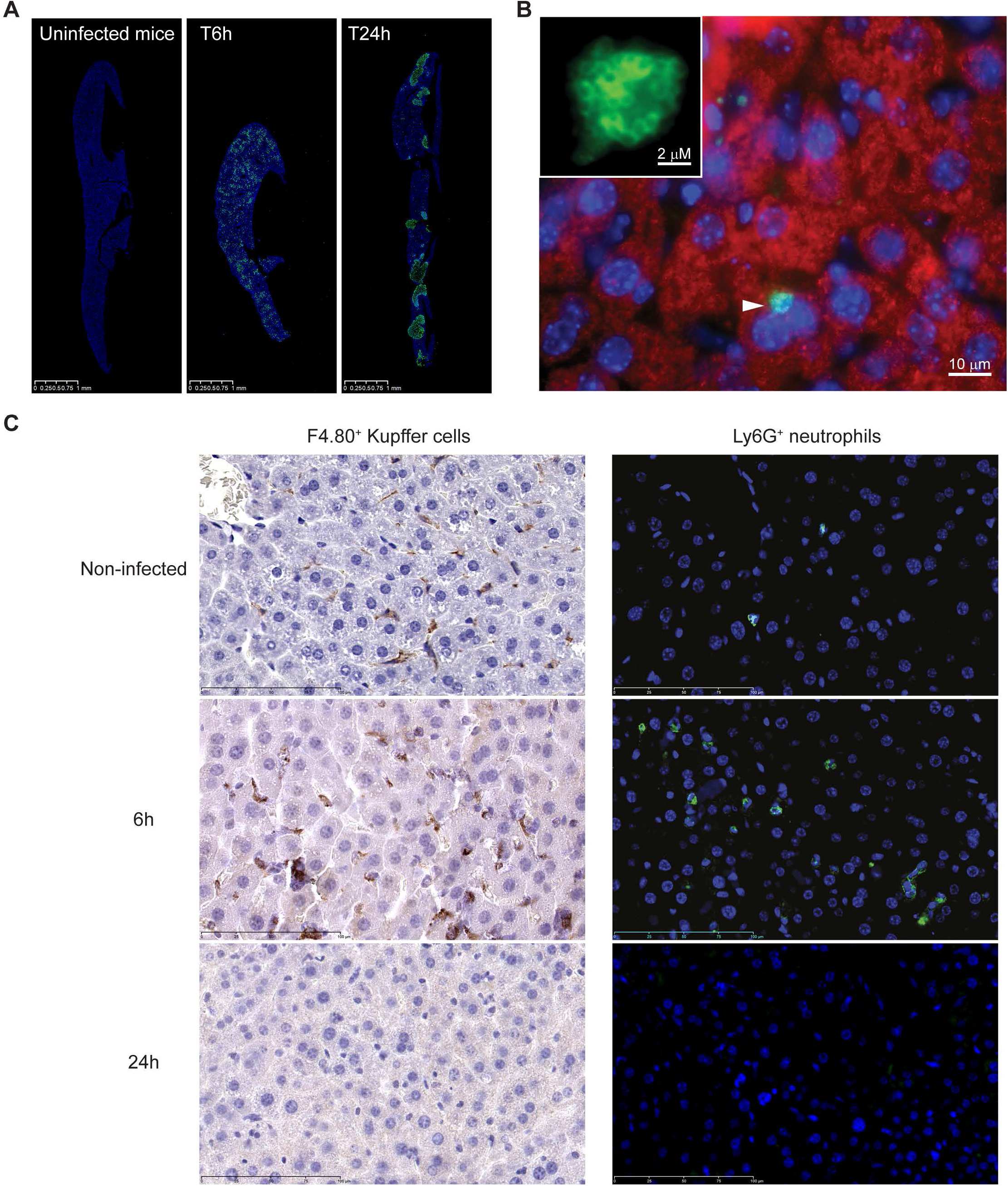
Formation of enterococcal clusters *in vivo* accompanies induction of the innate immune response. Female BALB/c mice were infected intravenously with 5×10^9^ CFUs of the OG1RF *lux* strain. **(A)** Representative images of liver histological sections after 6 and 24 h of infection are shown with noninfected control mice. Section of the largest liver lobe was labeled with anti-*Streptococcus* group D antiserum (*E. faecalis* in green) and Hoechst (nuclei in blue). **(B)** Intracellular *E. faecalis* clusters (green) were observed at 24 h in multinucleated (Hoechst, blue) claudin-1 expressing hepatocytes (red). A white arrowhead indicates an intracellular cluster in the hepatocyte, which is shown at a higher magnification (Bar: 2 µm). **(C)** Representative sections of the largest liver lobe labeled with hematoxylin and eosin and with an F4/80 antibody to detect Kupffer macrophages (brown cells, left panels), and with an anti-Ly6G antibody to stain neutrophils (green cells, right panels) and Hoechst (nuclei in blue).

Kupffer cells are resident liver macrophages that play a crucial role in the innate immune response and are responsible for the clearance of pathogens reaching the liver. The distribution of Kupffer cells was compared along the liver sections between noninfected mice and mice infected for 6 and 24 h with *E. faecalis*. Based on the surface area of noninfected mice, 35 F4/80^+^ macrophages/mm^2^ were observed. This number increased to about 53 macrophages/mm^2^ at 6 h pi. The number of macrophages decreased significantly to less than 5 macrophages/mm^2^ in mice infected for 24 h, showing an almost complete disappearance of the liver resident macrophages (Figure 3C). Although less abundant in the liver of control mice with 2 to 8 Ly6G^+^ neutrophils/mm^2^, the neutrophil population exhibited about a ten-fold increase, reaching 70 neutrophils/mm^2^ within the first 6 h of infection, before becoming almost undetectable 24 h pi. Together, these data show that the innate immune response to counteract *E. faecalis* infection is induced during the first hours of the infection, followed by a drastic depletion in macrophage and neutrophil cell density.

### *Is* E. faecalis *intracellular growth a common process?*

To investigate whether intracellular growth is a widespread process among *E. faecalis*, the behavior of two other human *E. faecalis* strains from distinct origins were tested in addition to our reference OG1RF strain, namely another clinical strain (JH2-2) and a probiotic strain (Symbioflor) in HepG2 cells. For all strains, the number of internalized bacteria were compared to the initial inoculum and an increase in the percentage of intracellular OG1RF and JH2-2 bacteria was observed (Figure 4A). In contrast, the percentage of intracellular bacteria did not significantly change for the Symbioflor strain, supporting that *E. faecalis* growth in hepatocytes is a strain-dependent process. Next, since a cluster of intracellular *E. faecalis* was observed in urothelial cells, as well as the presence of intracellular microcolonies ^18, 23^, *E. faecalis* growth was examined in two human kidney cancer cell lines. A704 and ACHN cells were used in a 48-h infection assay with the three *E. faecalis* strains. *E. faecalis* infection strongly differed between the two cell types. A704 kidney cells were more permissive for *E. faecalis* invasion and intracellular growth than ACHN cells, in which none of the strains grew (Figures 4B and 4C). In contrast to our observation during HepG2 infection, the level of intracellular Symbioflor strain increased in the A704 kidney cells 24 h pa. Although the same trend was observed for the OG1RF and JH2-2 strains, the intracellular level of each strain did not significantly change in kidney A704 cells. Notably, intracellular levels of OG1RF and JH2-2 decreased at 48 h, suggesting that the A704 cells may not be as permissive for the growth of these strains. To pursue this comparison, the percentage of kidney cells with intracellular OG1RF was determined according to the size of the clusters (Figure S4). Although large intracellular enterococcal clusters containing more than 10 enterococci were both detected in A704 and ACHN cells, the percentage of infected cells containing these large clusters remained low over time, compared to Huh7 hepatocytes (Figure 2B). Altogether, our results show that *E. faecalis* intracellular growth is not restricted to a specific cell type and may depend on a specific strain and cell-type combination.

**Fig 4.**
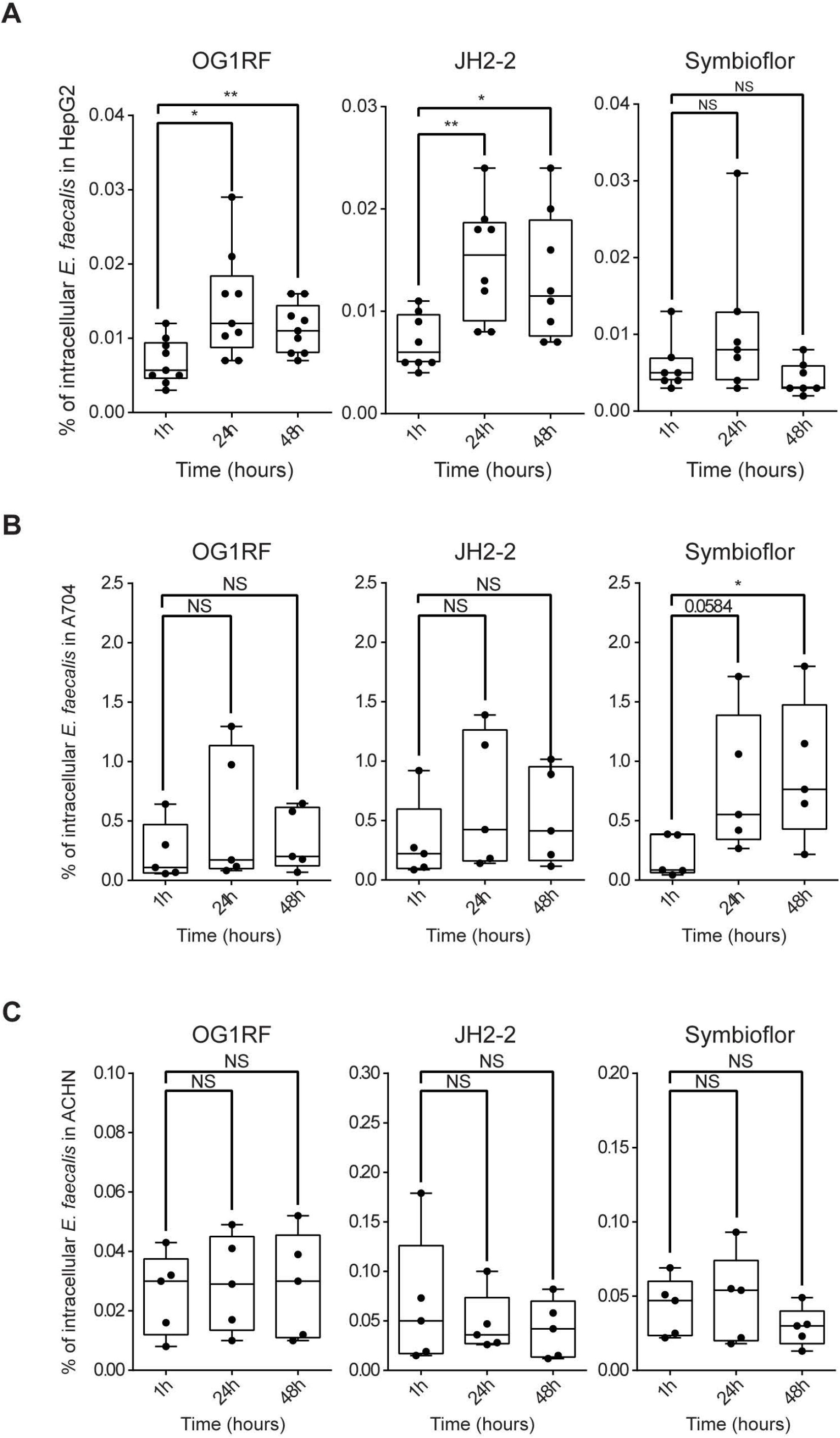
Intracellular growth of *E. faecalis* is a strain-and cell-type-dependent process. **(A)** Three *E. faecalis* strains (OG1RF, JH2-2, and Symbioflor) were used to infect HepG2 cells for 48 h. The percentage of intracellular bacteria was determined as the ratio of intracellular bacteria of the initial inoculum. All data are represented by a box-whiskers plot (Min to Max) of seven independent experiments. Each dot represents one independent experiment. The horizontal bar indicates the median value. Statistical analysis was performed using an unpaired Student’s t test. Asterisks indicate a p-value considered statistically significant (*, P < 0.05, **, P < 0.01), NS, non-significant difference. The three *E. faecalis* strains (OG1RF, JH2-2, and Symbioflor) were also used to infect A704 **(B)** and ACHN **(C)** kidney cells for 48 h. The percentage of intracellular bacteria was determined as the ratio of intracellular bacteria of the initial inoculum. All data are represented by a box-whisker plot (min to max) of five independent experiments. Each dot represents one independent experiment. The horizontal bar indicates the median value. Statistical analysis was performed using an unpaired Student’s t test. Asterisks indicate a p-value considered statistically significant (*, P < 0.05, **, P < 0.01), NS, non-significant difference.

## Discussion

*E. faecalis* is a unique opportunistic pathogen identified as contributing to liver damage in alcoholic liver disease ^36^. The present study demonstrated that *E. faecalis* replicates intracellularly in the liver. We highlighted that intracellular *E. faecalis* synthesizes peptidoglycan during the infection of hepatocytes and *E. faecalis* clusters form in two distinct hepatocyte cell lines, in primary mouse hepatocytes as well as *in vivo*. To our knowledge, this is the first demonstration of enterococcal intracellular division in mammalian cells. We also showed that induction of the liver innate immunity is followed by an almost disappearance of two major subsets, as the resident macrophages and the neutrophils, coinciding with the formation and spread of enterococcal foci. Finally, we showed that *E. faecalis* intracellular division can be extended to kidney cells. Overall, although all cell types may not be equally permissive for enterococcal growth, our findings indicate that this process is not restricted to a specific cell type. Indeed, during completion of this manuscript Tay *et al* reached similar conclusion that *E. faecalis* can survive and replicate after its internalization in keratinocytes ^37^.

The concept of an intracellular lifestyle has emerged quite recently for several opportunistic pathogens generally recognized as extracellular pathogens. O’Neill et al. (2016) provided the first direct evidence of group A *Streptococcus* replication inside human macrophages ^38^. Since then, intracellular replication of *Streptococcus pneumoniae* has also been observed in splenic macrophages ^39^. S*taphylococcus aureus*, which was historically regarded as a classical toxin-producing extracellular pathogen, is now widely accepted as a facultative intracellular pathogen ^40^. Very recently, Salcedo’s group described an intracellular niche for *Acinetobacter baumannii*, another nocosomial pathogen mainly described as an extracellular pathogen with restricted survival within cells ^41^. Some pathogenic fungi, such as *Blastomyces dermatitidis*, can also display a facultative intracellular lifestyle ^42^. Based on our findings and others ^18, 23^, *E. faecalis* can grow intracellularly and form microcolonies in hepatocytes, in kidney cells and in urothelial cells. In line with the contribution of *E. faecalis* to liver damage in alcoholic liver disease and its incidence with urinary tract infections, the liver, bladder, and kidneys are relevant target tissues. Future investigations on *E. faecalis* intracellular lifestyle will make sense, especially in light of the variety of organs or host sites targeted by *E. faecalis*.

Deciphering the cellular mechanisms and identifying the bacterial determinants supporting *E. faecalis* intracellular division in hepatocytes remains challenging. Several factors are known to be involved in *E. faecalis* stress tolerance and pathogenesis ^43^. Consistent with the generalist status and metabolic flexibility of *E. faecalis* isolates, intracellular growth of *E. faecalis* may be a strain-dependent mechanism. Moreover, the ability of the Symbioflor strain to grow intracellularly in kidney cells and not in hepatocytes supports that *E. faecalis* intracellular growth may require a specific strain and cell-type combination. Among several mechanisms, hijacking the host endocytic and autophagy pathways is a common strategy for intracellular pathogens ^44^. *E. faecalis* is able to survive for up to 72 h within macrophages ^25^. Zou and Shankar (2016) showed that *E. faecalis* can delay lysosomal fusion of the enterococcal-containing compartment in macrophages. They found two types of enterococcal populations with some *E. faecalis* surrounded by single membrane vacuoles and some that had lost their vacuolar compartment, suggesting that the latter may escape and reside in the cytoplasm ^24^. Autophagy is a conserved process in which cytoplasmic components are targeted to the lysosomes for degradation. While *E. faecalis* entry into epithelial cells is a well-admitted process, the epithelial cell intrinsic mechanisms that detects and targets intracellular *E. faecalis* has begun to be explored. Hooper’s team showed that autophagy in intestinal epithelial cells was activated by *E. faecalis*, which is entrapped in double-membrane autophagosomes ^27^. Their results also suggested that *E. faecalis* reaches the cytosol during the infectious process. Tay *et al*. showed that once internalized into keratinocytes via macropinocytosis in single membrane-bound compartments, some intracellular *E. faecalis* are detected in early and late endosomes and proposed that intracellular replication occurs within late endosomes until a threshold is reached and some bacteria are released into the cytosol ^37^. In line with their findings, we observed very few *E. faecalis* in Rab5- or EEA1-positive compartments during the first hours of the infection (data not shown). Moreover, at later time points, enterococcal clusters were not localized into acidic compartments. Finally, the almost complete disappearance of intracellular *E. faecalis* in hepatocytes exposed to amoxicillin, which diffuses through cell membranes and penetrates the cytoplasm, further supports that at least one enterococcal population is located in the cytosol, where it divides.

The liver plays a major role in the clearance and response to commensal bacteria translocating from the gastrointestinal tract and to enteropathogens, such as *E. coli* and *Listeria monocytogenes*. This protective role is mostly mediated by resident Kupffer macrophages. The latter participate in the innate immune response at several levels by clearing bacteria, secreting soluble inflammatory mediators, and/or physically interacting with effectors of other cell types ^45^. Rapid recruitment of neutrophils is an additional important line of defense, particularly for the rapid clearance of *E. faecalis* and *E. faecium* ^46, 47^. Accordingly, severely ill patients with hematologic malignancies and deep neutropenia were at an increased risk of developing enterococcal infections. Here, a strong recruitment of neutrophils and an increase in Kupffer macrophages were observed in the livers of mice infected with *E. faecalis*, followed by an almost disappearance of both cell types, coinciding with the formation and spread of enterococcal foci. We hypothesize that the oxidative burst generated in Kupffer cells and neutrophils leads to the depletion of innate cells in the liver, creating favorable conditions for the formation of intracellular enterococcal clusters. In their most severe form, liver diseases are associated with a high risk of mortality, and treatment options are often limited. Identifying factors that contribute to the onset and progression of liver injury is necessary to improve the management of patients with liver diseases. *E. faecalis* translocation to the liver leads to detrimental inflammation for the host in several rodent models by extracellular bacteria ^14, 16, 48^. The intracellular location of *E. faecalis* in hepatocytes may be a protective niche against immune detection and may favor the establishment of *E. faecalis* in the liver. If extracellular *E. faecalis* has been shown to mediate inflammation in the liver upon intestinal translocation in a mouse model ^14, 16, 48^, the possibility of intracellular *E. faecalis* within hepatocytes contributing to liver disorders has not been considered thus far. Further studies will help to determine how this novel intracellular lifestyle may contribute to liver diseases and possibly other diseases.

## Materials and methods

### Bacterial strains

*E. faecalis* strains OG1RF ^49^, JH2-2 ^50^, and Symbioflor (a gift from Dr. E. Domann Institute of Medical Microbiology, University of Giessen, Germany) were cultured in brain heart infusion (BHI) at 37°C without aeration. GFP-expressing OG1RF from the pMV158-GFP plasmid was cultured in BHI with 4 µg/ml tetracycline ^51^. *E. faecalis* strain OG1RF, expressing the *luxABCDE* (*lux*) operon from *Photorhabdus luminescens*, was a gift from Dr. D. Lechardeur (Micalis Institute, INRAE, Centre de Recherche Ile de France—Jouy-en-Josas - Antony) ^52^ and was cultured in BHI with 20 µg/ml erythromycin.

### Cell lines

The human hepatocellular carcinoma Huh7 cell line (CLS 300156) was cultured in Dulbecco’s modified eagle medium (DMEM, Gibco) with glutamax supplemented with 10% FBS. The human hepatocellular carcinoma HepG2 cell line (ATCC HB-8065) and the human kidney adenocarcinoma A-704 cell line (ECACC 93020513) were grown in minimum essential medium (MEM, Gibco) with glutamax supplemented with 10% FBS, 0.1 mM non-essential amino acids, and 1 mM sodium pyruvate. The human kidney adenocarcinoma ACHN cell line (ECACC 88100508) was grown in MEM with glutamax with 10% FBS and 0.1 mM non-essential amino acids. All cell lines were cultured at 37°C in a 5% CO_2_ atmosphere.

### Cell infection

Two or four days before infection, cells were seeded in triplicate in 24-well plates or on glass coverslips for immunofluorescence analysis. Prior to infection, cells were washed once with PBS and incubated in serum-free medium for 2 h. *E. faecalis* strains were grown until bacteria reached the mid-exponential phase. Bacteria were harvested, washed twice in phosphate-buffered saline (PBS), and resuspended in medium without serum to be used at a multiplicity of infection (MOI) of 50. Infection was synchronized by 1 min centrifugation at 1000 g. After 3 h of contact, cells were washed 5 times with PBS, and an antibiotic cocktail was added to kill extracellular bacteria. A first antibiotic cocktail (150 µg/ml gentamicin and 10 µg/ml vancomycin) was added for 24 h and then replaced by another antibiotic cocktail (37.5 µg/ml gentamicin and 5 µg/ml vancomycin) for the rest of infection. When indicated, amoxicillin was added to the antibiotic cocktail (125 µg/ml and diluted at 50 µg/ml after 24 h of infection). The efficiency of the antibiotic cocktails was controlled by the absence of viable colonies after plating of the cell supernatants. When required, cells were lysed using cold distilled water for 10 min at 4°C to enumerate intracellular bacteria on BHI agar plates or processed for immunofluorescence as described below. The percentage of intracellular bacteria was determined as the ratio of intracellular bacteria of the initial inoculum.

### Peptidoglycan labelling

The Huh7 hepatocytes were seeded in a cell culture µ-dish (Clinisciences, ibidi 81156) four days before infection and then infected with GFP-expressing OG1RF *E. faecalis* as described in the section on cell infection. After 12 h of infection, 1 mM orange-red TAMRA-based fluorescent D-amino acid (RADA, Tocris, 6649) was added to the µ-dish. After 38 h of infection, cells were washed 5 times in Hanks’ balanced salt solution (HBSS) 1X and fixed in 4% PFA for 20 min at room temperature. The Hoechst stain (Sigma B2261, 5 µg/ml) was used to stain DNA. As a control, remaining antibiotic-killed extracellular enterococci were detected using the rabbit anti-*Enterococcus* antiserum (diluted 1:1000) and a goat anti-rabbit-Alexa Fluor 647-conjugated secondary antibody (ThermoFisher Scientific, A-21244 diluted 1:200), as described above.

### Extracellular/intracellular bacterial staining

At each time point, cells were washed three times in PBS-Ca-Mg buffer (Gibco, DPBS, calcium, magnesium) and fixed with 4% paraformaldehyde (PFA) in PBS-Ca-Mg for 20 min. Fixed cells were washed twice with PBS-Ca-Mg and blocked using 5% BSA in PBS-Ca-Mg for 20 min. To discriminate intracellular bacteria from the remaining antibiotic-killed extracellular bacteria, double staining was performed where extracellular bacteria were stained prior to cell permeabilization, as previously described ^34^. Briefly, for extracellular bacteria staining, the infected cells were incubated with a rabbit anti-*Streptococcus* group D antiserum (BD Diagnostics, Le Pont de Claix, France) diluted at 1:1000 in 2% BSA in PBS-Ca-Mg for 1 h, washed 3 times, and incubated with a secondary goat anti-rabbit IgG antibody conjugated with Cy3 (Amersham Biosciences 1:400 in BSA 2% in PBS-Ca-Mg) for 1 h. After cell permeabilization, the rabbit anti-*Enterococcus* antiserum was used for 1 h prior to a secondary goat anti-rabbit IgG antibody conjugated with Alexa Fluor 488 diluted at 1:500 in BSA 2% in PBS-Ca-Mg for 1 h to label intracellular bacteria. The Hoechst stain (Sigma B2261, 5 µg/ml) was used to stain DNA. Samples were mounted on glass coverslips and analyzed with a fluorescent microscope (Carl Zeiss Axiovert 135, AxioObserver.Z1, KEYENCE BZ-X710). Images were acquired with a 40× or 100× oil immersion objective using a Zeiss Axiocam 506 camera. Image quantification analysis was performed using Zen software (Carl Zeiss) and Image J software. At least 80 images (taken with 40×) and 50 images (taken with 100×) were quantified in total from three independent experiments.

### Ethics statement

All animals were housed under specific pathogen-free conditions in our local animal facility (IERP, INRAE, Jouy-en-Josas). Mice were fed irradiated food and autoclaved water ad libitum, in line with animal welfare guidelines. The animal house was maintained on a 12-h light/dark cycle. Animal experiments were approved by the local ethics committee, the COMETHEA (“Comité d’Ethique en Expérimentation Animale du Centre INRAe de Jouy-en-Josas et AgroParisTech”), under registration number 19-08 and by the French Ministry of Higher Education and Research (APAFIS #20380-2019060315249683 v1) and were performed in accordance with European directive 2010/63/EU.

### Primary mouse hepatocyte isolation and culture

Primary mouse hepatocytes (PMH) were isolated from 8–10-week-old female C57BL/6 mice. PMH were isolated by collagenase perfusion of the liver, as previously described ^53^. Briefly, mice were anesthetized with xylazine (10 mg/kg IP) and ketamine (100 mg/kg IP) and subject to a mid-line laparotomy. The inferior vena cava was perfused with a 0.05% collagenase solution (collagenase from *Clostridium histolyticum*, Sigma C5138). The portal vein was sectioned, and the solution allowed to flow through the liver. Upon collagenase digestion, hepatic cells were removed by mechanical dissociation, filtered through a sterile 70 μm cell strainer (BD Falcon), and washed twice by centrifugation at 300 g for 4 min. After a filtration step through a sterile 40 μm cell strainer (BD Falcon), cells were resuspended in serum-containing culture medium (DMEM Gibco, 10% fetal bovine serum, 1% penicillin– streptomycin, and 100 µg/mL Fungizone). Cell count and viability were assessed by trypan blue exclusion. Cells (500,000 cells/well) were seeded in 6-well collagen-coated plates for 6 h at 37°C in a 5% CO2 atmosphere. After complete adhesion of the hepatocytes and washes to remove the dead cells, PMH were cultured in hepatocyte culture medium (William’s E medium, GlutaMAX™ Supplement, Gibco™ 32551020; 100 U/ml penicillin/streptomycin, Sigma P4333; 0.5 µg/ml Fungizone antimycotic B, Gibco 15290018; 4 µg/ml insulin, Sigma I0516; 0.1 % bovine serum albumin, Sigma A8412 and 25 nM dexamethasone, Sigma D2915) at 37°C in a 5% CO2 atmosphere for 4–6 days before infection.

### Mouse infection and luminescence imaging

Experiments were conducted on 9- to 10-week-old adult female BALB/cByJRj mice (Janvier Labs). All animals were adapted to the environment of the local animal facilities (IERP, INRAE, Jouy-en-Josas) for one week prior to the study. *E. faecalis* strain OG1RF, expressing the *lux* operon, was collected by centrifugation 1 h after bacteria had reached the stationary phase. Bacterial cells were washed twice with PBS buffer and stored at −80°C. Mice were infected intravenously in the retro-orbital vein with 5×10^9^ CFUs. Serial dilutions of the inoculum were also transferred to plates as a control for determining inoculated *E. faecalis* numbers. Bioluminescent enterococci were imaged from mice under isoflurane anesthesia using the *In Vivo* Imaging System (IVIS spectrum BL, PerkinElmer) equipped with Living Image software (version 4.7.3, PerkinElmer) as reported previously ^52^. When required, mice were sacrificed by cervical dislocation. Bioluminescence images of the mice were acquired with a 23.1 cm field of view (FOV). For liver lobes, the FOV value was 13.5. Photon emission was measured as radiance (photons per second per square centimeter per steradian, p.s^-1^.cm^-2^.sr^-1^). All luminescence images were adjusted on the same color scale and corrected (final pixel size: binning 4; pixel size smoothing: 5×5).

### Histology and immunostaining

Livers were fixed overnight in 4% paraformaldehyde and then embedded in paraffin. Immunohistochemistry was performed on paraffin sections (5 μm) using antibody F4-80 (Bio-rad, France; diluted 1:100). Sections were incubated overnight at 4°C, washed and incubated with an appropriate biotinylated secondary antibody for 1 h at room temperature, and then with streptavidin-HRP complex followed by 3,3-diaminobenzidine or 3-amino-9-ethylcarbazole detection (LSAB kit, Dako France). Sections were then counterstained with hematoxylin. For immunofluorescence staining, sections were labelled with antibodies, rabbit anti-*Enterococcus* serum (diluted 1:2000), mouse monoclonal anti-claudin 1 (Clinisciences, sc-166338 diluted 1:100), and mouse anti-Ly6G (Biolegend, France, clone A8; diluted 1:50), followed by staining with appropriate secondary antibodies, Alexa FluorTM 488 (Invitrogen, ThermoFisher Scientific A-11008 diluted 1:250) or DyLight550 (ThermoFisher Scientific 84540, diluted 1:250). Nuclei were stained with Hoechst 33342 (molecular probes). All sections were scanned using a NanoZoomer 2.0-RS digital slide scanner (Hamamatsu, Japan). Images were digitally captured from the scanned slides using NDP.view2 software (Hamamatsu). Sections labeled with claudin-1 were acquired with a 100× oil immersion objective using a Zeiss Axiocam 506 camera. Images were processed with Zen software (Carl Zeiss).

### Statistical analyses

Statistical analyses were performed with Prism 6 (GraphPad Software). An unpaired two-tailed Student’s t-test was used to compare the means of the two groups. For multiple comparisons, data analysis was performed using the Kruskal-Wallis test with Dunnett’s post-hoc test for comparison of means relative to the mean of a control group.

## Acknowledgments

We are grateful to the IERP unit, INRAE (Infectiology of fishes and rodent facility, doi: 10.15454/1.5572427140471238E12) and thank J. Rivière and M. Vilotte from the histology facility of UMR 1313 GABI, 78350, Jouy-en-Josas, France. We thank D. Lecharcheur for the gift of the *E. faecalis* OG1RF strain expressing the *luxABCDE* (*lux*) operon from *Photorhabdus luminescens*. We are very grateful to D. Deschamps for technical advice and assistance with IVIS experiments. We thank the Emerg’in platform for access to IVIS-Spectrum BL, which was financed by the Region Ile de France (DIMOneHealth).

## Availability of data

All data generated or analyzed during this study are included in this published article and its supplementary information files.

## Authors’ contributions

NN, ABD, GL, AL, FMB, and CA performed experiments and analyzed data. HB, AMC, PS and CA were involved on the conceptualization. PS and CA supervised and wrote the manuscript; all authors reviewed and approved the final manuscript.

## SUPPLEMENTARY FIGURE LEGENDS

**Supp. Fig S1.** Representative micrographs of extracellular *E. faecalis* upon Huh7 infection with the GFP-expressing *E. faecalis* OG1RF and incubated with the orange-red TAMRA-based fluorescent D-amino acid (RADA). The image is an overlay of the phase contrast, extracellular *E. faecalis* (pink channel), and nuclei (blue channel). The scale bar corresponds to 5 µm. Absence of RADA signal on antibiotic-killed extracellular *E. faecalis* is shown at a higher magnification below (Bar: 1 μm).

**Supp. Fig S2. Quantification of the formation of *E. faecalis* intracellular clusters in Huh7 hepatocytes**. Exponentially growing *E. faecalis* OG1RF cultures were used to infect Huh7 cells for 48 h. Percentage of cells exhibiting at least one cluster **(A)**, as well as the number of enterococci (triangle symbol) by clusters, **(B)** are shown. For each time point, at least 600 cells were examined at high magnification (objective 100×) from three independent experiments. Asterisks indicate statistical significance compared with the 30 min condition using the Kruskal-Wallis one-way ANOVA followed by Dunnett’s test for multiple comparisons (*, P < 0.05, **, P < 0.01; ***, P < 0.001). All data are represented by a box-whiskers plot (min to max) of at least four independent experiments. Each dot represents one independent experiment. The horizontal bar indicates the median value.

**Supp. Fig S3. *In vivo* imaging of *E. faecalis* infection**. Female BALB/c mice were infected intravenously with 5×10^9^ CFUs of the OG1RF *lux* strain. At 6 and 24 h post-infection, anesthetized mice were imaged using the IVIS 200 system. Representative animals and livers from two independent experiments are shown. Photon emission was measured as radiance (photons per second per square centimeter per steradian, p.s^-1^.cm^-2^.sr^-1^). Luminescence images from mice were adjusted for the color scale at minimum or Min=8×10^4^ and maximum or Max=1.5×10^6^ p.s^-1^.cm^-2^ sr^-1^. Luminescence images from the liver were adjusted at Min= 1×10^5^ and maximum or Max=2.5×10^6^ p.s^-1^.cm^-2^.sr^-1^. Rainbow images show the relative level of luminescence ranging from low (blue) to high (red).

**Supp. Fig S4. Formation of intracellular *E. faecalis* clusters in kidney cells**. Quantification of the percentage of cells according to the number of enterococci within the intracellular cluster in A704 and ACHN cells. For each time point, at least 2,500 cells were examined at low magnification (objective 40×) from three independent experiments.

