## Supplementary figures and images for "The unforeseen intracellular lifestyle of *Enterococcus faecalis* in hepatocytes"

### Supplemental Figures

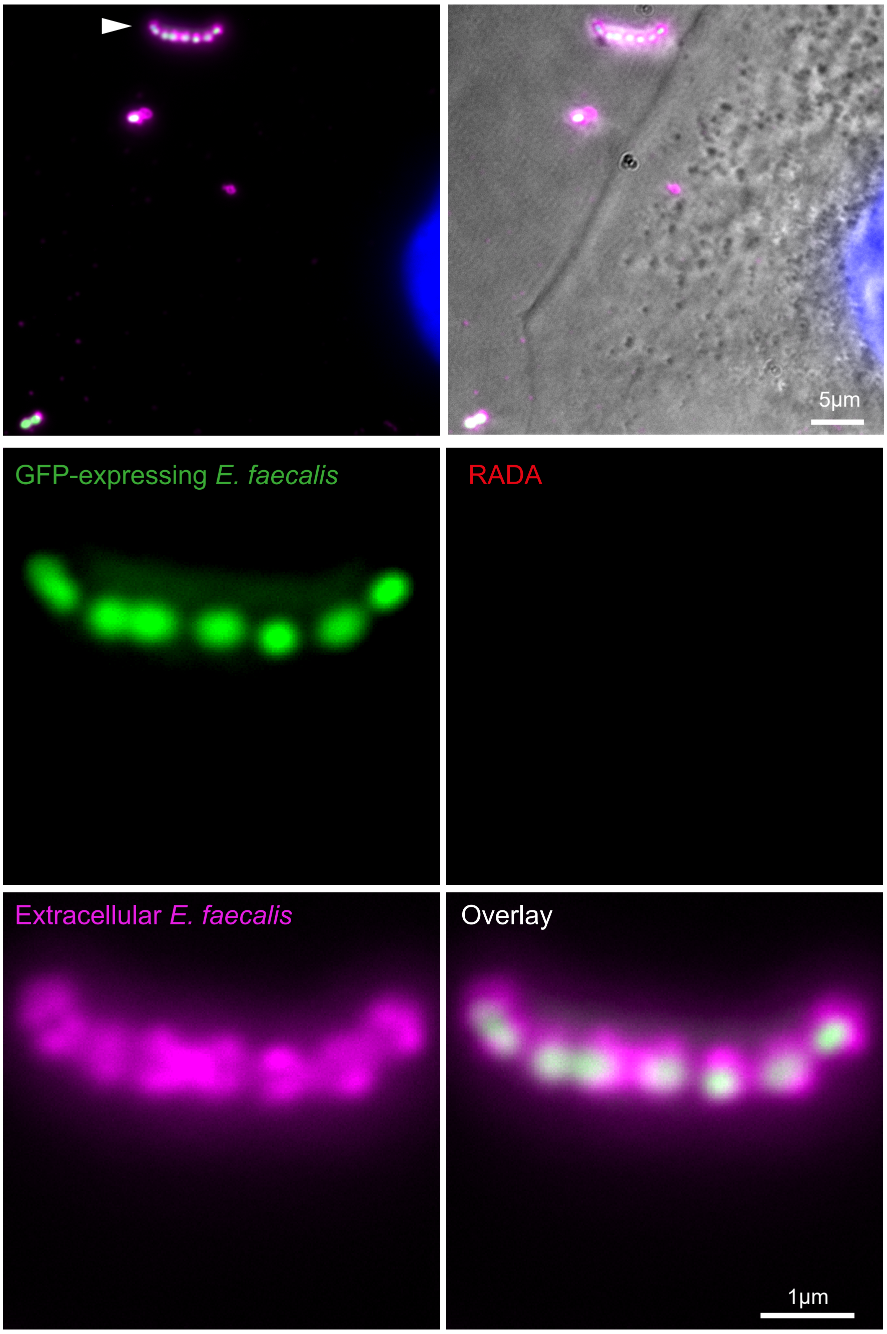

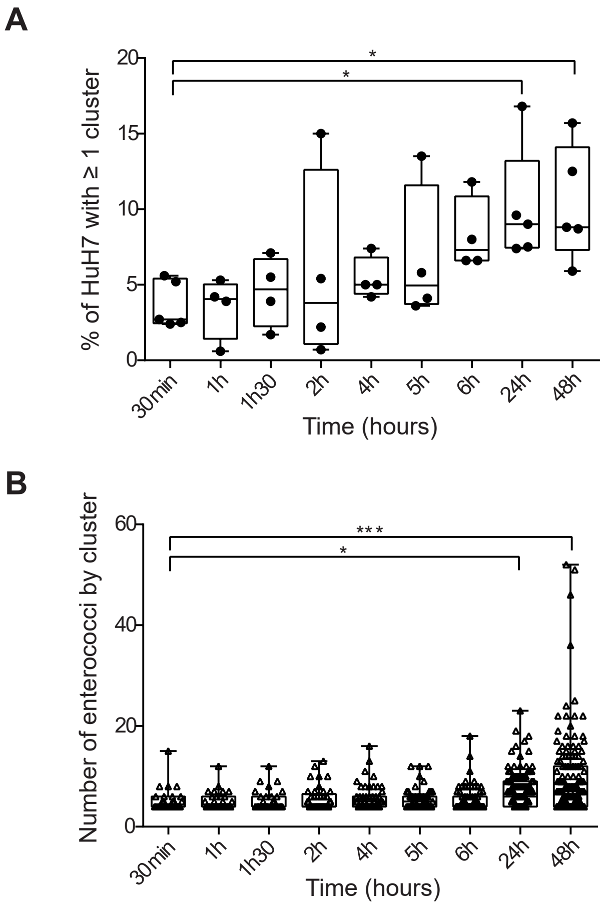

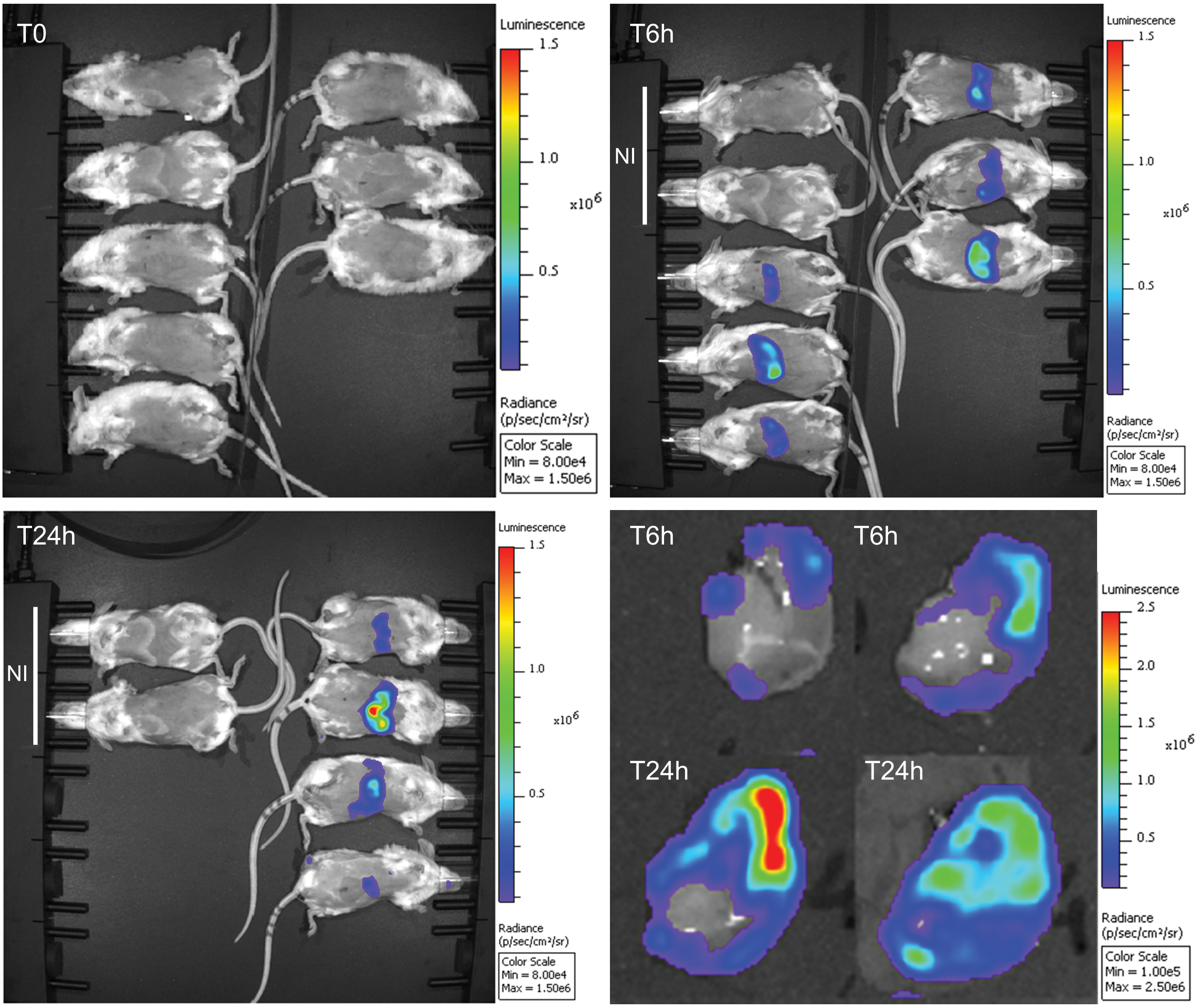

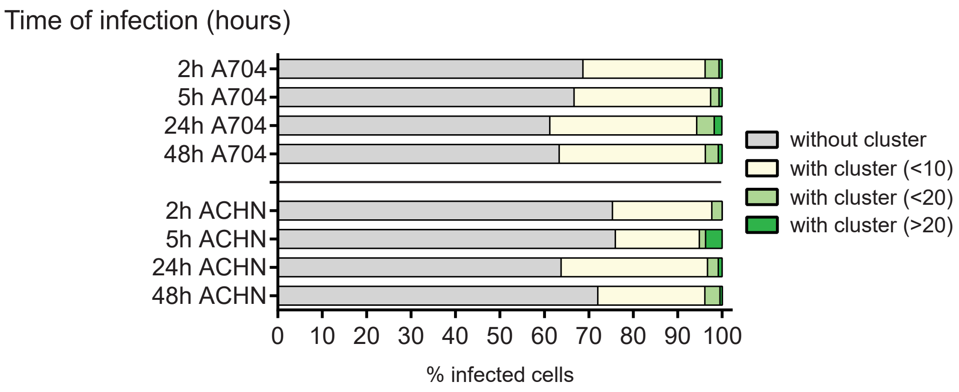
